# Bridging human and plant adaptations for climate resilience

**DOI:** 10.64898/2026.02.20.706989

**Authors:** Nicola Favretto, Heng Lim Tan, Gemma Brain, Daphne Ezer

## Abstract

Climate change is reshaping agriculture through both gradual shifts and increasingly unpredictable extremes. Plants cope using developmental plasticity and bet-hedging, but it is unclear how these biological strategies align with the ways farmers perceive and respond to climate risks. This study investigates: (1) whether farmers understand climate change as incremental trends or recurrent shocks, (2) how their adaptations parallel plant plasticity and bet-hedging, and (3) under which climate scenarios these adaptations best support yield stability.
We combined qualitative research and modelling by conducting fifty semi-structured interviews with farmers, agricultural associations and public administrators across three climatically distinct Italian regions, and by developing an agent-based stochastic simulation that represents farmer-like plasticity (delayed sowing) and bet-hedging (staggered sowing) under drought and flood scenarios.
Farmers described climate change as both gradual transformation and intensifying volatility. Their adaptive responses – adjusting calendars, switching crops and diversifying production – closely aligned with plant strategies, though articulated in practical rather than scientific terms. Simulation results showed that plasticity enhanced yields under systematic shifts in conditions, whereas bet-hedging reduced losses in highly variable climates characterised by frequent transitions between extremes.
Together, the qualitative and modelling findings demonstrate that plant and farmer adaptation logics converge in complementary ways. Plasticity supports performance under gradual change, while bet-hedging buffers unpredictability. These insights highlight the potential for co-designed tools that link plant traits, farmer decision-making and ecological risk, strengthening climate-resilient agricultural planning and improving communication between farmers, breeders and plant scientists.

**Societal Impact Statement:** Climate change is transforming agriculture through both gradual shifts and increasingly unpredictable extremes, challenging farmers’ ability to protect crops and livelihoods. This study brings together farmer experiences and plant adaptation strategies to explore how people and plants respond to similar climate pressures. By showing that farmers’ practices mirror plant plasticity and bet-hedging, our findings highlight opportunities to design climate-resilient agriculture that aligns biological traits with real-world decision-making. This work can inform plant breeders, extension services and policymakers seeking to support farmers through clearer communication, better risk-management tools and more adaptable crop varieties, ultimately strengthening resilience in food systems.

## 1. INTRODUCTION

Global temperatures are increasing at an unprecedented pace, accompanied by shifts in precipitation patterns and more frequent extreme weather events, which are altering agro-climatic zones and influencing crop choices (Muluneh, 2021). Projections suggest that some regions, including Northern Europe, Canada, Siberia, and Eastern Russia, may become more suitable for agriculture (Santos et al., 2020), while others – such as Southern and Eastern Europe, sub-Saharan Africa, South Asia, and Central America – face heightened risks to yields from recurrent floods and droughts. Evidence from global assessments (IPCC, 2022), machine learning projections under Representative Concentration Pathways (RCPs) scenarios of meteorological drought (Shiru et al., 2020), and modelling of historical weather-induced crop failures (Goulart et al. 2021) consistently support these trends.

Plants have evolved multiple strategies to cope with unpredictable environmental fluctuation (Starrfelt and Kokko, 2012). One of the main strategies is developmental plasticity, whereby plants permanently alter traits such as leaf form, root depth, or flowering time to match prevailing stresses (Gawinowski et al., 2025). For example, leaves may thicken under strong light and heat, or elongate petioles under high temperatures to improve cooling (Scheepens et al., 2018). Plasticity maximises performance under current conditions, but can limit flexibility if the environment later changes (Botero et al., 2015). A second strategy is diversified bet-hedging, which spreads risk across offspring (Simons, 2011). In many species, seeds within the same pod germinate at different times: some sprout immediately, others remain dormant (Abley et al., 2024). While delaying germination reduces short-term success in favourable years, it ensures survival after shocks such as drought (Gianella et al., 2021). Together, plasticity and bet-hedging offer plants complementary tools for coping with both predictable and unpredictable stresses (Joschinski and Bonte, 2020).

Just as plants adapt to current environmental stresses or adjust their behaviour in anticipation of future ones, farmers also respond to increasingly erratic climate patterns and extreme weather events. From a socio-environmental management perspective, these responses aim to maintain yields and minimise economic losses. Farmer adaptations include a variety of nature-based, land-based, and ecosystem-based approaches – such as watershed management, afforestation, and agroforestry – as well as livelihood diversification, including crop diversification and changes to planting schedules (Favretto and Stringer, 2024).

Bridging plant science with human decision-making reveals a dual challenge rooted in two key research gaps. First, crop and yield models often fail to account for both plant adaptive strategies and human behavioural responses (Albanito et al., 2022). This lack of socio-economic integration and disconnection between plant responses and farmer practices severely limits the ability to translate biological insights into practical guidance for end users. Second, many stakeholders, particularly farmers, have limited familiarity with biological concepts that could enhance their adaptive decision-making, and often face difficulties in identifying reliable, accessible sources of useful information (Rust et al., 2022). This gap highlights a broader systemic issue: the absence of a two-way knowledge exchange mechanism that connects biological research with real-world agricultural decision-making.

This paper bridges human and plant adaptations to climate change through a transdisciplinary case study that examines how farmers and plants converge on similar strategies of survival. It combines farmers’ perceptions of climate change and lived experiences of adaptation in Italy with simulations of plant plasticity and bet-hedging under alternative climate scenarios. Three guiding questions structure the analysis at the interface of plant and social sciences:

1. How do farmers in Italy perceive climate change – primarily as gradual shifts in conditions or as more frequent extreme weather events – and what consequences do they report for crop performance?”
2. How are farmers adapting to perceived climate change impacts, and how closely do their practices overlap with plant adaptations such as plasticity and bet-hedging?
3. How might farmer-like plasticity and bet-hedging adaptations influence crop yields under varying climate scenarios?

Together, these questions link empirical evidence with theoretical insights from plant biology and simulation modelling. Farmers’ accounts provide grounded insights that inform the design and interpretation of the simulations, while the modelling results help contextualise the adaptations farmers describe. This integrated approach highlights how social science perspectives on perceptions and practices can enrich experimental models of plasticity and bet-hedging, generating more socially relevant decision-support tools. It offers a novel and globally relevant perspective on the adaptive logics that shape resilience under climate uncertainty, with timely contributions to support farmers and decision-makers confronting complex risks to food systems and livelihoods.

## 2. MATERIALS AND METHODS

### 2.1 Social science

Italy was selected as the study area for farmer interviews due to its exposure to climate-related challenges that mirror broader European and global trends. Rising temperatures, declining water availability, and increasingly severe weather events – such as droughts and floods – are already having profound effects on agriculture. In 2022 alone, rice and wheat yields fell by 30% due to drought (Coldiretti, 2022; MASE, 2023). To explore farmers’ adaptation responses to these climatic pressures, research was conducted in three contrasting Italian regions: Aosta Valley, Basilicata, and Sardinia. These were selected to reflect a diversity of climate vulnerabilities (mountain, hill, arid, and semi-arid zones), population sizes, land use characteristics, and the economic emphasis on agriculture, capturing a representative cross-section of regional conditions within Italy (ISTAT, 2019a; 2019b, 2019c and 2023).

Qualitative data were collected through fieldwork conducted between October 2023 and January 2024. Ethical approval was granted by the University of York, UK (Ref: DEGERC/Res/23062023/1), and all participants provided informed consent. Regional stakeholders were initially identified through national-level contacts, enabling purposive sampling which was then expanded via snowball sampling (Parker, 2019). Interviews focused on farmers’ perceptions of climate impacts and adaptations. Sampling continued until theoretical saturation was reached, following principles aligned with a grounded theory approach (Glaser and Strauss, 1999). In total, 50 semi-structured interviews were conducted: 17 with individual farmers or farmer cooperatives, 7 with representatives from national farmers’ associations (Coldiretti, Confagricoltura, CIA, and Copagri), which together represent over 3 million agricultural enterprises across Italy, and 26 with public administrators from the environment, agriculture and water sectors (see Table 1).

**Table 1.** Interview sample by region and stakeholder type.

|  | Aosta Valley | Sardinia | Basilicata | Total |
| --- | --- | --- | --- | --- |
| Individual farmers and cooperatives | 4 | 5 | 8 | **17** |
| Farmers associations | 1 | 3 | 3 | **7** |
| Public administration (Environment, Agriculture and Water) | 9 | 8 | 9 | **26** |
| Total |  |  |  | **50** |

All interviews were audio-recorded with consent, transcribed, and analysed using thematic analysis in NVivo 14. The analysis followed four key steps: data import and organisation, coding in line with the research questions, identification of cross-cutting themes, and selection of excerpts. A total of 57 coded excerpts were retained for presentation in the results (Nowell et al., 2017) (see Supporting Information, Interview excerpts S1 and S2).

### 2.2. Simulation

An agent-based stochastic simulation was developed in R (4.3.2) to model how different weather patterns impact the yield benefits from plasticity and bet-hedging-related farmer adaptations. All of the quantities are unitless.

#### Weather patterns

Weather patterns were defined as a sequence of 10 weather events (normal, drought, flood). We generated weather patterns in two different ways:

i. Random. Each weather event was randomly and independently selected. The probability of selecting drought or flood were equal and the probability of an adverse weather event varied between 0% and 66%. This represents an increasing incidence of extreme weather events.
ii. Markov chain. Weather events were generated by a Markov chain, so that the weather in one time point is more likely to be consistent with the previous time point. The Markov chain was initialised with 80% normal weather, 10% drought and 10% flood on the first day. The transition rates to other weather conditions varied between 0% and 66%. This represents weather “trends”, where the current weather is indicative of future weather.

#### Plants

When a seed is sown, a new plant is initialised with an age and stress level of 0. Each plant had an Aging Rate (AR) drawn from a normal distribution (mean=1, stdev=0.5). The developmental stage of the plant is determined by its internal age (age<3 is a seed, 3<=age<10 is vegetative, 10<=age<12 is post-bolting, and age>=12 is ready to harvest, again these are unitless parameters). Within each time period, the plants’ age and stress level is adjusted based on the weather pattern assigned to that time period and the current developmental stage of the plant, as per Table 2.

**Table 2.**
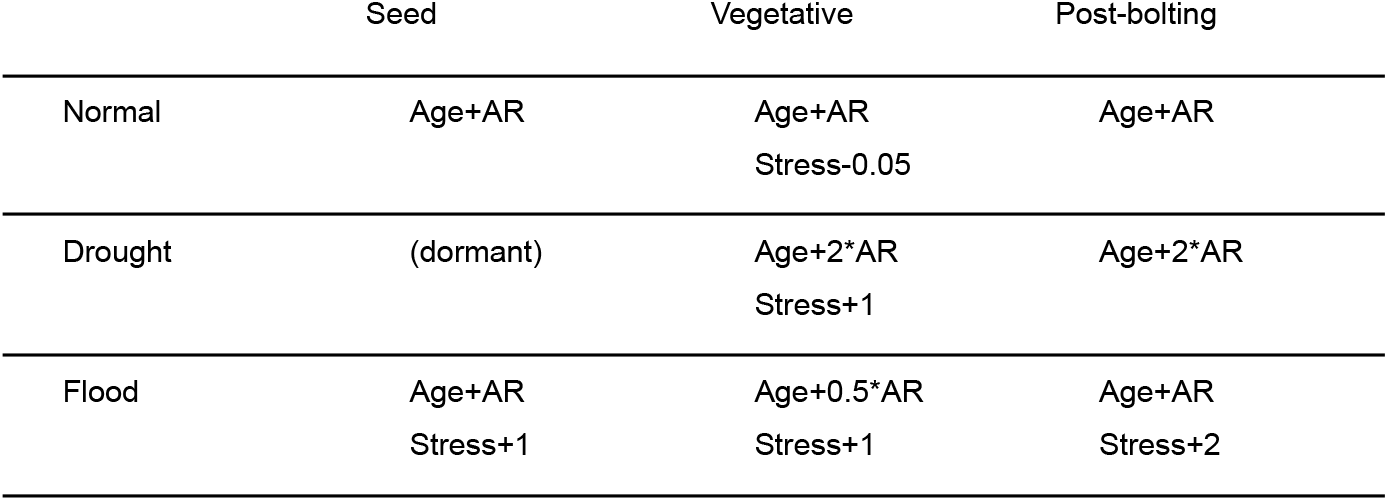
Interview sample by region and stakeholder type.

These parameters were selected to reflect certain principles in plant biology: under droughts, in most cereals, seed germination is delayed, vegetative plants have more rapid transition to flowering and senescence, and vegetative plants wilt, reducing overall yields (Havrlentová et al., 2021). Under flooding, germination can occur, as long as conditions are aerobic (Guglielminetti et al., 1995), but plants are under continuous greater risk of rot (Schafer and Kotanen, 2003), especially those that are further along in their development (Mills et al., 2020).

Within each time period, a certain number of plants will die, based on their current stress levels and developmental stage. For each plant, a number is drawn from a normal distribution with a mean of that plant’s current stress level and a standard deviation of 0.2. Vegetative phased plants that draw a number over 3 or post-bolting plants that draw a number over 4 will die and be removed from the simulation. These parameters were selected so that death was a rare event among plants grown in normal conditions.

#### Farmers

In the simulation, farmers can choose one of three alternatives: business-as-usual, plasticity, or bet-hedging. Under business-as-usual, farmers sow all their seeds on the first day of the simulation. Under plasticity, farmers are willing to wait for favourable conditions: they will either sow their seeds on the first day without floods or droughts or after 10 time periods (unitless), whichever comes first. Under bet-hedging, farmers diversify their sowing schedules by sowing half their field on the first day of the simulation and then waiting for 5 time periods before sowing the remainder of the seeds. In all cases, they harvest their crops 10 time periods after their seeds are sown. Only plants that are ready to harvest and have 10 or fewer stress points can provide yield. Every plant provides a yield that is linearly penalised by its stress level (0.1*stress).

As inherent in any stochastic simulation of this nature, many of the parameters were artificially set. However, the entire simulation is available here (stressedplants/PlasticityBetHedging: Simulation of how farmer decisions influence yield), enabling a dynamic exploration of how the parameter space influences modelling outcomes.

Figure 1 maps the research questions to their respective data sources, highlighting how social and plant science findings are brought together.

**Figure 1.**
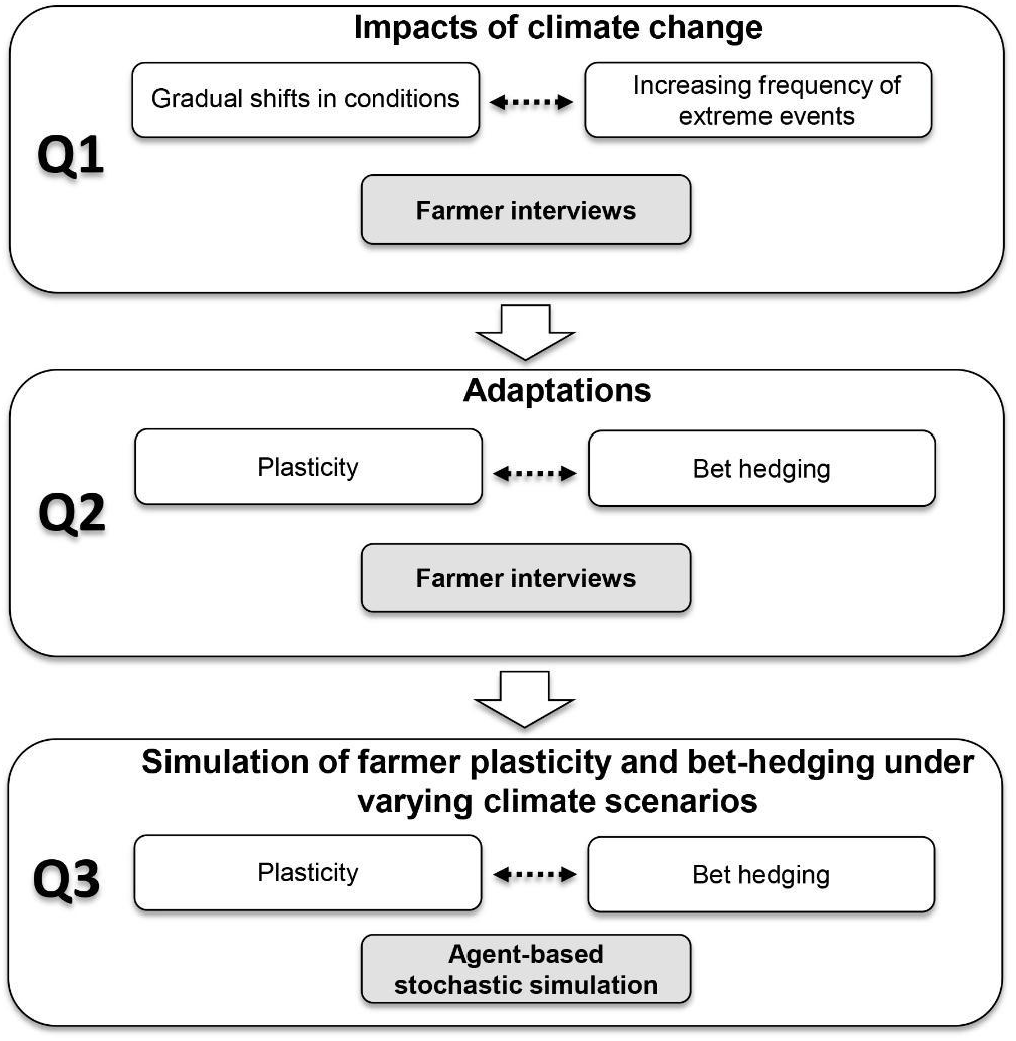
Connections between research questions and data sources. Q refers to research question numbers, each linked to categories of climate impacts (gradual shifts vs. increasing frequency), adaptations (plasticity vs. bet-hedging), and simulation approaches (plasticity vs. bet-hedging). Shaded boxes indicate data sources. Dotted arrows represent reciprocal linkages between types of impacts or adaptations, while vertical arrows illustrate the sequential progression.

## 3. RESULTS AND DISCUSSION

### 3.1 Farmers’ perceptions of climate change impacts

Farmers across Italian regions report profound climate-driven changes, pointing to both gradual shifts in conditions and an increasing frequency of extreme events. Their accounts mirror wider Mediterranean trends, where rising temperatures (Barcikowska et al., 2020; Hochman et al., 2022), altered rainfall (Hochman et al., 2019), and recurrent extremes (Perkins-Kirkpatrick and Lewis, 2020) are reshaping agriculture. The testimonies highlight how these dynamics are simultaneously experienced as incremental seasonal adjustments and as disruptive shocks that expose new vulnerabilities.

#### Gradual shifts in seasonal conditions

Farmers describe how slow-moving changes in weather patterns undermine established routines. Water stress increasingly affects spring–summer crops such as maize and wheat when irrigation is unavailable (Guerriero et al., 2024). Drought delays sowing and disrupts germination, leading one public administrator in Basilicata to remark: “Spring has shifted by a few months” (public administrator, Basilicata). Here, both plants and farmers exhibit plasticity – delayed germination in crops, and delayed sowing by farmers – reflecting coupled adjustments under gradual but predictable conditions (Giménez-Benavides et al., 2011). Several farmers note that combinations of gradual changes amplify challenges. A recurring pattern of heavier winter or spring rains followed by prolonged summer droughts heightens water stress, alters growing seasons, and reduces yields and quality. In the Aosta Valley, one farmer observed: “Agri-systems are undergoing changes in the length of their phenological cycles, productivity, and shifts in traditional cultivation areas” (farmer, Aosta Valley). This illustrates a dual plasticity: plants shorten or extend phenological cycles, while farmers shift cultivation zones. Similarly, in Sardinia, a farmer explained: “Due to summer drought and heavy rains in May, the olive’s growth process has shifted. The fruit ripening phase has been delayed” (farmer, Sardinia). This aligns with evidence of Mediterranean olive cultivars adjusting their developmental programmes to climate pressures (Didevarasl et al., 2025).

#### Interactions with extreme events

Gradual trends also interact with extreme events to magnify risks. In Basilicata, warmer winters (a slow trend) trigger catastrophic pest outbreaks: “During the Christmas period, we had highs of 15°C… they compromised the entire harvest” (farmer, Basilicata). In Sardinia, one farmer recalled: “In the past, when cold came, plants would go into dormancy… but now they don’t stop, they keep growing” (farmer, Sardinia). Continuous growth, once advantageous, now exposes crops to sudden freezes and long-term economic losses. Above-average winter temperatures induce early budding, leaving crops vulnerable to late frosts: “One April frost could destroy everything” (farmer, Sardinia). Persistent rains similarly foster disease in vines and olives, with a public administrator reporting that “in 2023… losses exceeded 90%” (public administrator, Basilicata). Such effects are rarely confined to a single season, as stress in one year diminishes production for years to follow. These accounts highlight a central tension: while plasticity allows adaptation to current conditions, such adjustments can become maladaptive under heightened volatility, echoing ecological debates about the limits of plasticity (Botero et al., 2015).

#### Normalisation of extremes

A striking theme is the normalisation of extreme conditions. Farmers stress that recurrent shocks are no longer perceived as rare events but as part of the baseline climate. As one farmer put it: “We’re getting used to these extreme conditions. Heat now arrives earlier, and this winter was constantly warm” (farmer, Basilicata). This perception resonates with observations across the Eastern Mediterranean and Middle East, where extremes are shifting toward longer-term climatic patterns (Zittis et al., 2022). Such accounts blur the conventional dichotomy between gradual change and episodic shocks, reframing extremes as the “new normal”.

#### Absent from the discourse: trait heterogeneity

While farmers consistently describe plastic responses in both crops and farming practices, they rarely mention trait heterogeneity, a hallmark of diversified bet-hedging (Gianella et al., 2021). Variability in traits such as germination timing or developmental rates enhances resilience at the population level by ensuring that some individuals survive adverse events (Pinceel et al., 2021). Yet uniformity has long been favoured in modern crops, making this strategy largely invisible to farmers. Its absence from their discourse suggests a blind spot in adaptation thinking, with implications for breeding programmes and on-farm resilience.

Taken together, farmers’ perceptions reveal three important insights. First, they capture the interplay between gradual shifts and extreme events, showing how the former heightens vulnerability to the latter. Second, they highlight parallels between plant and farmer plasticity, where both biological and human systems recalibrate to changing conditions, though often with limited effectiveness under volatility. Third, they expose the normalisation of extremes, resonating with global calls for a redefinition of what constitutes “baseline” climate. The silence around trait heterogeneity is equally telling. Farmers’ emphasis on plasticity but neglect of bet-hedging adaptations underscores a disconnection between plant science and farming practice. Bridging this gap could inform both crop breeding and decision-support tools, strengthening resilience in the face of climate uncertainty.

### 3.2 Farmers’ adaptations overlapping with plasticity and bet-hedging

In response to climatic uncertainty, farmers adopt a wide range of adaptations that reflect elements of plasticity (adjusting practices in real time to prevailing conditions) and bet-hedging (spreading risk to prepare for future volatility). As one farmer explained: “Agriculture is plastic; once farmers realise something can’t be done anymore due to changing conditions, they change crops and irrigation periods, seeking a crop that can still ensure profitability” (farmer, Basilicata).

#### Real-time adjustments (plasticity)

Precision agriculture technologies are increasingly deployed to monitor and respond to environmental changes. In Sardinia, farmers “monitor critical phases… which prevents damage” (farmer, Sardinia) while in the Aosta Valley “satellite and drone-based surveys, relying on NDVI or infrared indices” (farmer, Aosta Valley) guide irrigation timing. Farmers also adjust the timing of sowing, fertilisation, irrigation, and harvesting. In Sardinia, cropping calendars have shifted: “The irrigation season used to start in April and end in November; now we have spring cycles for herbaceous plants or autumn–winter cycles for lettuces” (farmer, Sardinia). In Basilicata, wheat sowing has moved from November to December or January to avoid losses from excessive rainfall: “Those who sowed in November lost their crops; seeds rotted in the soil and didn’t germinate” (farmer, Basilicata). Fertilisation schedules are likewise restructured, with applications delayed until rainfall is likely: “If it doesn’t rain, we risk doing more harm than good” (farmer, Basilicata).

#### Crop and varietal change (plasticity and bet-hedging overlap)

Farmers also alter the crops they grow when formerly reliable varieties become unviable. In Sardinia: “It may get replaced within the same farm or by neighbouring farms; crops are evolving into slightly different ones” (farmer, Sardinia). A fruit grower in Basilicata reported sourcing new clementine varieties from Spain: “Every year we attend their workshops; where they present the latest varieties that enable us to keep producing under changing climatic conditions” (farmer, Basilicata). Agronomists similarly note a reorientation toward species not dependent on cold winters: “We are reorienting cultivation toward crops that don’t require winter cold to complete their cycle” (agronomist, Basilicata).

Such practices highlight a continuum between plasticity and bet-hedging. Complete replacement of crops in response to observed change is a plastic adaptation, mirroring shifts in olive cultivation across latitudinal ranges (Morea, 2021). For example, in drier zones of Basilicata, olives increasingly replace citrus: “It’s a good response in arid regions and performs well under rising temperatures” (olive oil producers association, Basilicata). Conversely, diversification into multiple crops spreads risk and more closely aligns with bet-hedging.

#### Diversification (bet-hedging)

Crop diversification has become widespread, and has been supported by EU greening payments since 2015 (Runge et al., 2022). A national survey showed that by 2020, 88% of Italian farms had diversified, with reduced risks compared to monocultures (Fabri et al., 2024). Regional data confirm increasing diversity across southern Italy (Monteleone et al., 2018). Farmers echo this logic: diversification ensures at least some crops survive adverse conditions, mirroring biological adaptations where populations stagger germination or flowering times to spread risk: “There’s a move toward diversifying production in some areas, especially where summer crops are no longer viable” (Farmer, Basilicata). In practice, however, such measures are often adopted primarily to comply with EU Common Agricultural Policy (CAP) requirements rather than out of a conscious bet-hedging rationale: “We find ourselves rotating crops in line with the requirements imposed by the CAP eco-schemes” (farmer, Sardinia).

#### Protective infrastructure (risk reduction beyond bet-hedging)

Farmers also invest in protective measures such as anti-hail nets and windbreaks: “We use protection systems that serve both as hail nets and to shield plants from intense solar radiation and low humidity” (farmer, Sardinia). These interventions create microclimates that reduce exposure to volatility. While not strictly plasticity or bet-hedging, they reflect the same anticipatory logic of minimising exposure to climatic extremes.

Farmers’ lived experiences reveal how adaptations align with plant logics of survival. Real-time adjustments in calendars and irrigation mirror plant plasticity, where development is tuned to immediate conditions. Diversification and varietal change reflect elements of bet-hedging, reducing the likelihood of total failure under future uncertainty. Protective infrastructure represents a complementary layer of risk buffering. The ambiguity between plasticity and bet-hedging – whether a crop change is a flexible adjustment or a diversification hedge – suggests these adaptations are not dichotomous but exist on a continuum. Farmers, like plants, combine both approaches, underscoring the value of viewing human adaptation through the dual lens of plasticity and bet-hedging.

### 3.3 Modelling plasticity- and bet-hedging-type farmer adaptations to climate impacts

As shown by the interview data, Italian farmers perceive both gradual shifts and greater volatility in seasonal weather patterns, with particular concern for the increasing frequency and duration of floods and droughts. Among the most frequently discussed adaptations is the adjustment of seasonal activities, especially sowing dates. Farmers’ practices align closely with plant adaptations of plasticity (real-time adjustment) and bet-hedging (diversification to spread risk). To model how farmer agricultural adaptations that mirror plant plasticity and bet-hedging influence crop yields under varying climate scenarios, we developed a stochastic agent-based simulation of these adaptations under varying frequencies and durations of drought and flood events (Figure 2).

**Figure 2.**
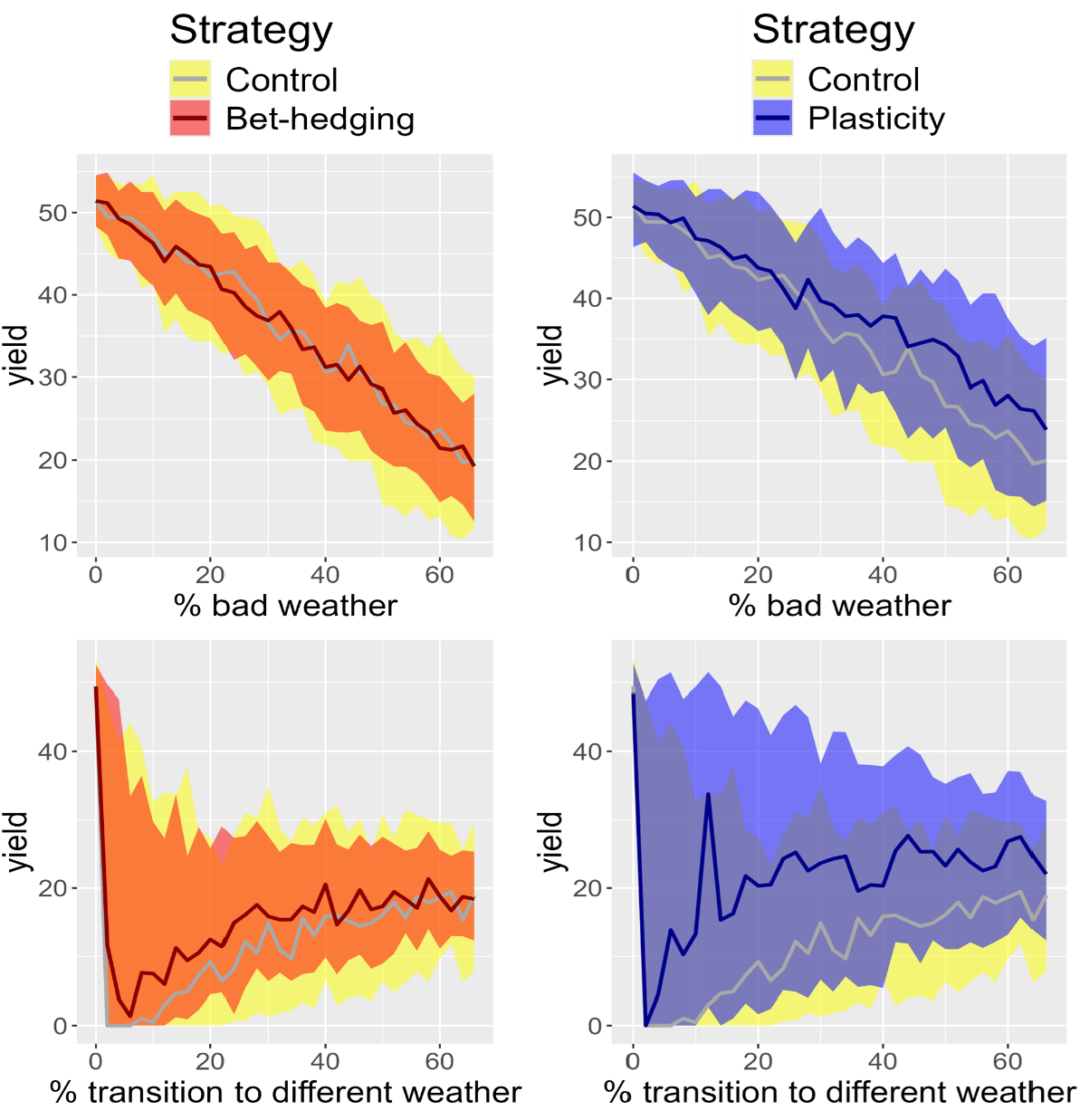
Stochastic simulation results of farmer bet-hedging and plasticity adaptations under increasing frequency of floods/droughts (% bad weather) and decreasing consistency of patterns (% transition to different weather). Lines represent medians; coloured ribbons show interquartile ranges. N=100 for each simulated scenario.

Previous research suggests that bet-hedging adaptations – such as staggered germination or variable growth rates – can increase population resilience under drought (McCarthy et al., 2025). Translating this logic, we modelled farmer bet-hedging as staggering sowing times across fields, a common practice in many farming communities (Derbile, 2013) and a proxy for species diversification. While not explicitly raised in our Italian interviews, this mechanism allows us to test the potential benefits of diversification. Results confirm that farmer bet-hedging reduces yield volatility, consistent with studies showing that crop diversification lowers risk in Italian farms (Rosa et al., 2019). Our results suggest that diversification through sowing-time staggering may therefore reduce risk.

Plasticity was modelled as a willingness to delay sowing until favourable conditions emerge. The simulations show that plasticity substantially improves yields under longer-duration droughts and floods. However, under highly volatile conditions with frequent transitions between extremes, delayed sowing is less effective. This aligns with ecological findings that plant plasticity is most advantageous under gradual shifts in climate regimes, while its benefits diminish under high-frequency variability (Alpert and Simms, 2002).

Overall, our model demonstrates that plasticity and bet-hedging each confer advantages under different types of climatic stress. Plasticity is most effective when adverse conditions persist for extended periods, whereas bet-hedging reduces losses under unpredictable variability. Together, they provide complementary adaptations for coping with climate change.

## 4. CONCLUSION

This study shows how farmer perceptions and practices echo the adaptive strategies long observed in plants. Farmers in Italy experience climate change as both incremental shifts and intensifying extremes, and their responses — adjusting calendars, switching crops, diversifying production — mirror plant plasticity and bet-hedging, even without explicit reference to biological concepts. Our simulations reinforce these parallels: plasticity is most advantageous under gradual shifts, whereas bet-hedging buffers volatility when conditions fluctuate unpredictably. Importantly, the distinction between plasticity and bet-hedging blurs in practice. Farmers combine short-term flexibility, diversification, and protective measures, suggesting adaptation operates along a continuum rather than through discrete strategies. This perspective opens space for more integrative adaptation frameworks linking plant traits, farmer decision-making, and resilience planning. The findings highlight opportunities for co-designed tools and breeding approaches that align plant adaptive capacities with farmer priorities and risk perceptions. Strengthening two-way communication between farmers, breeders and plant scientists could support more sustainable seed selection, diversification pathways, and climate-resilient agricultural systems. By drawing together ecological theory and lived experience, this work underscores the shared adaptive logics that connect plants, people and the landscapes they depend upon

## ACKNOWLEDGMENTS

This research was funded by the York Environmental Sustainability Institute (YESI) Discipline Hopping Scheme at the University of York, by the University of York Department of Environment and Geography and the BBSRC (BB/S506795/1) and EPSRC (EP/W524657/1). We thank Prof. Lindsay Stringer for reviewing the manuscript.

## AUTHOR CONTRIBUTIONS

N.F. designed and led the project, conducted interviews, and wrote the manuscript. D.E. co-designed the project, developed the model, revised the manuscript, and supervised H.L.T. and G.B. H.L.T. and G.B. contributed to bioscience framing and manuscript review.

## DATA AVAILABILITY STATEMENT

This study did not generate new unique materials/reagents. The qualitative interview data generated in this study cannot be shared via online repositories in order to protect participant anonymity. Summarised excerpts are provided in the Supporting Information. Simulation code used for model development is available here: https://github.com/stressedplants/PlasticityBetHedging.

## REFERENCES

Abley, K., Goswami, R., Locke, J.C.W. (2024). Bet-hedging and variability in plant development: seed germination and beyond. Phil Trans R Soc B 379(1900): 20230048. 10.1098/rstb.2023.0048.

Albanito, F., et al. (2022). How modelers model: the overlooked social and human dimensions in model intercomparison studies. Environ Sci Technol 56(18): 13485–13498. 10.1021/acs.est.2c02023.

Alpert, P., Simms, E.L. (2002). The relative advantages of plasticity and fixity in different environments: when is it good for a plant to adjust? Evol Ecol 16: 285–297. 10.1023/A:1019684612767.

Barcikowska, M.J., Kapnick, S.B., Krishnamurty, L. et al. (2020). Changes in the future summer Mediterranean climate: contribution of teleconnections and local factors. Earth Syst Dynam 11: 161–181. 10.5194/esd-11-161-2020.

Botero, C.A., Weissing, F.J., Wright, J. et al. (2015). Evolutionary tipping points in the capacity to adapt to environmental change. PNAS 112: 184–189. 10.1073/pnas.1408589111.

Coldiretti. 2022. Clima: 6 mld di danni dalla peggiore siccità da 500 anni. Available at: https://www.coldiretti.it/meteo_clima/clima-6-mld-di-danni-dalla-peggiore-siccita-da-500-anni. [Accessed on 05/05/2024].

Derbile, E.K. (2013). Reducing vulnerability of rain-fed agriculture to drought through indigenous knowledge systems in north-eastern Ghana. Int J Climate Change Strateg Manag 5(1): 71–94. 10.1108/17568691311299372.

Didevarasl, A., Costa-Saura, J.M., Spano, D. et al. (2025). The phenological phases of early and mid-late budbreak olive cultivars in a changing future climate over the Euro-Mediterranean region. Eur J Agron 168: 127658. 10.1016/j.eja.2025.127658.

Fabri, C., Vermeulen, S., Van Passel, S. et al. (2024). Crop diversification and the effect of weather shocks on Italian farmers’ income and income risk. J Agric Econ 75. 10.1111/1477-9552.12610.

Favretto, N., Stringer, L.C. (2024). Climate resilient development in vulnerable geographies. Mitig Adapt Strateg Glob Change 29: 90. 10.1007/s11027-024-10187-5.

Gawinowski, M., Chenu, K., Deswarte, J.C. et al. (2025). Plant plasticity in the face of climate change – CO2 offsetting effects to warming and water deficit in wheat: a review. Environ Exp Bot 232: 106113. 10.1016/j.envexpbot.2025.106113.

Gianella, M., Bradford, K.J., Guzzon, F. (2021). Ecological, (epi)genetic and physiological aspects of bet-hedging in angiosperms. Plant Reprod 34: 21–36. 10.1007/s00497-020-00402-z.

Giménez-Benavides, L., García-Camacho, R., Iriondo, J. et al. (2011). Selection on flowering time in Mediterranean high-mountain plants under global warming. Evol Ecol 25: 777–794. 10.1007/s10682-010-9440-z.

Glaser, B., Strauss, A. (1999). The discovery of grounded theory: strategies for qualitative research. Aldine Transaction: Piscataway, NJ, USA. ISBN 0-202-30260-1.

Goulart, H.M.D., van der Wiel, K., Folberth, C. et al. (2021). Storylines of weather-induced crop failure events under climate change. Earth Syst Dynam 12: 1503–1527. 10.5194/esd-12-1503-2021.

Guerriero, V., Scorzini, A.R., Di Lena, B. et al. (2024). Measuring variation of crop production vulnerability to climate fluctuations over time: the case of wheat in Abruzzo, Italy. Sustainability 16(15): 6462. 10.3390/su16156462.

Guglielminetti, L., Yamaguchi, J., Perata, P. et al. (1995). Amylolytic activities in cereal seeds under aerobic and anaerobic conditions. Plant Physiol 109(3): 1069–1076. 10.1104/pp.109.3.1069.

Havrlentová, M., Kraic, J., Gregusová, V., Kovácsová, B. (2021). Drought stress in cereals – a review. Agriculture 67(2). 10.2478/agri-2021-0005.

Hochman, A., Kunin, P., Alpert, P. et al. (2019). Weather regimes and analogues downscaling of seasonal precipitation for the 21st century: a case study over Israel. Int J Climatol. 40: 2062–2077. 10.1002/joc.6318.

Hochman, A., Marra, F., Messori, G. et al. (2022). Extreme weather and societal impacts in the eastern Mediterranean. Earth Syst Dynam 13: 749–777. 10.5194/esd-13-749-2022.

IPCC. (2022). Annex II: Glossary. In: Climate Change 2022: Impacts, Adaptation and Vulnerability. Contribution of Working Group II to the Sixth Assessment Report of the IPCC. Cambridge University Press: Cambridge, UK and New York, USA. 10.1017/9781009325844.029.

ISTAT. (2019a). Censimento permanente delle imprese: Report Basilicata 2019. National Institute of Statistics. https://www.istat.it/wp-content/uploads/2021/03/CPUE_BASILICATA.pdf.

ISTAT. (2019b). Censimento permanente delle imprese: Report Sardegna 2019. National Institute of Statistics. https://www.istat.it/wp-content/uploads/2021/03/CPUE_SARDEGNA.pdf.

ISTAT. (2019c). Dati statistici per il territorio Regione Valle d’Aosta. National Institute of Statistics. https://www.istat.it/it/files/2020/05/02_Valle-dAosta_Scheda.pdf.

ISTAT. (2023). Censimento della popolazione: dati regionali - Anno 2023. National Institute of Statistics. https://www.istat.it/comunicato-territoriale/censimento-della-popolazione-dati-regionali-anno-2023/.

Joschinski, J., Bonte, D. (2020). Transgenerational plasticity and bet-hedging: a framework for reaction norm evolution. Front Ecol Evol 8: 517183. 10.3389/fevo.2020.517183.

MASE. 2023. Piano Nazionale di Adattamento ai Cambiamenti Climatici. Ministero dell’Ambiente e della Sicurezza Energetica. Available at: https://www.mase.gov.it/sites/default/files/PNACC_DOCUMENTO_DI_PIANO.pdf. [Accessed : 01/05/2025].

McCarthy, K., Pook, H., Redmond, E. et al. (2025). ELF3 controls trait heterogeneity by tuning the rate of maturation in Arabidopsis and barley. bioRxiv 2025.03.24.644900. 10.1101/2025.03.24.644900.

Mills, K.B., Madden, L.V., Paul, P.A. (2020). Quantifying the effects of temperature and relative humidity on the development of wheat blast incited by the Lolium pathotype of Magnaporthe oryzae. Plant Dis 104: 2622–2633. 10.1094/PDIS-12-19-2709-RE.

Monteleone, M., Cammerino, A.R.B., Libutti, A. (2018). Agricultural “greening” and cropland diversification trends: potential contribution of agroenergy crops in Capitanata (South Italy). Land Use Policy 70: 591–600. 10.1016/j.landusepol.2017.10.038.

Morea, D. (2021). Moving toward the north? The spatial shift of olive groves in Italy. Zemědělská Ekonomika 67: 129–135. 10.17221/467/2020-AGRICECON.

Muluneh, M.G. (2021). Impact of climate change on biodiversity and food security: a global perspective. Agric Food Secur 10: 36. 10.1186/s40066-021-00318-5.

Nowell, L.S., Norris, J.M., White, D.E. et al. (2017). Thematic analysis: striving to meet the trustworthiness criteria. Int J Qual Methods 16: 1. 10.1177/1609406917733847.

Parker, C., Scott, S., Geddes, A. (2019). Snowball sampling. In Atkinson, P., Delamont, S., Cernat, A., Sakshaug, J.W., Williams, R.A. (eds.), SAGE Research Methods Foundations. 10.4135/9781526421036831710.

Perkins-Kirkpatrick, S.E., Lewis, S.C. (2020). Increasing trends in regional heatwaves. Nat. Commun. 11: 3357. 10.1038/s41467-020-16970-7.

Pinceel, T., Buschke, F., Geerts, A., et al. (2021). An empirical confirmation of diversified bet hedging as a survival strategy in unpredictably varying environments. Ecology 102, e03496. 10.1002/ecy.3496.

Rosa, F., Taverna, M., Nassivera, F., et al. (2019). Farm/crop portfolio simulations under variable risk: a case study from Italy. Agric. Econ. 7, 8. 10.1186/s40100-019-0127-7.

Runge, T., Latacz-Lohmann, U., Schaller, L., et al. (2022). Implementation of eco-schemes in fifteen European Union member states. EuroChoices 21, 19–27. 10.1111/1746-692X.12352.

Rust, N.A., Stankovics, P., Jarvis, R.M. et al. (2022) Have farmers had enough of experts? Environ. Manage. 69: 31–44. 10.1007/s00267-021-01546-y.

Santos, J.A., Fraga, H., Malheiro, A.C. et al. (2020). A review of the potential climate change impacts and adaptation options for European viticulture. App. Sci. 10:3092. 10.3390/app10093092.

Schafer, M., Kotanen, P.M. (2003). The influence of soil moisture on losses of buried seeds to fungi. Acta Oecologica 24: 255–263. 10.1016/j.actao.2003.09.001.

Scheepens, J.F., Deng, Y., Bossdorf, O. (2018). Phenotypic plasticity in response to temperature fluctuations is genetically variable, and relates to climatic variability of origin, in Arabidopsis thaliana. AoB Plants 10, ply043. 10.1093/aobpla/ply043.

Shiru, M.S., Shahid, S., Dewan, A. at al. (2020). Projection of meteorological droughts in Nigeria during growing seasons under climate change scenarios. Sci. Rep. 10, 10107. 10.1038/s41598-020-67146-8.

Simons, A.M. (2011). Modes of response to environmental change and the elusive empirical evidence for bet hedging. Proc. R. Soc. B 278: 1601–1609. 10.1098/rspb.2011.0176.

Starrfelt, J., Kokko, H. (2012). Bet-hedging: a triple trade-off between means, variances and correlations. Biol Rev Camb Philos Soc 87(3): 742–755. 10.1111/j.1469-185X.2012.00225.x.

Zittis, G., Almazroui, M., Alpert, P., et al. (2022). Climate change and weather extremes in the Eastern Mediterranean and Middle East. Rev. Geophys. 60, e2021RG000762. 10.1029/2021RG000762.

